# Survival protection mechanisms and genetic variability induction after stress: two sides of the same Hsp70 coin

**DOI:** 10.1101/423210

**Authors:** Ugo Cappucci, Fabrizia Noro, Assunta Maria Casale, Laura Fanti, Maria Berloco, Angela Alessandra Alagia, Luigi Grassi, Loredana Le Pera, Lucia Piacentini, Sergio Pimpinelli

**Author notes:** these authors contributed equally to this work (Ugo Cappucci, Fabrizia Noro).

## Abstract

Previous studies have shown that heat shock stress may increase transcription levels and, in some cases, also the transposition of certain transposable elements (TEs) in *Drosophila* and other organisms. Other studies have also demonstrated that heat shock chaperones as Hsp90 and Hop are involved in repressing transposon’s activity in *Drosophila melanogaster* by their involvement in crucial steps of the biogenesis of Piwi-interacting RNAs (piRNAs), the largest class of germline-enriched small non-coding RNA implicated in the epigenetic silencing of TEs. However, a satisfying picture of how many chaperones and their respective functional roles could be involved in repressing transposons in germ cell is still unknown. Here we show that in *Drosophila* heat shock activates transposon′s expression at post-transcriptional level by disrupting a repressive chaperone complex by a decisive role of the stress-inducible chaperone Hsp70. We found that stress-induced transposons activation is triggered by an interaction of Hsp70 with the Hsc70-Hsp90 complex and other factors all involved in piRNA biogenesis in both ovaries and testes. Such interaction induces a displacement of all such factors to the lysosomes resulting in a functional collapse of piRNA biogenesis. In support of a significant role of Hsp70 in transposon activation after stress, we found that the expression under normal conditions of Hsp70 in transgenic flies increases the amount of transposon transcripts and displaces the components of chaperon machinery outside the nuage as observed after heat shock. So that, our results demonstrate that heat shock stress is capable to increase the expression of transposons at post-transcriptional level by affecting piRNA biogenesis through the action of the inducible chaperone Hsp70. We think that such mechanism proposes relevant evolutionary implications. In presence of drastic environmental changes, Hsp70 plays a key dual role in increasing both the survival probability of individuals and the genetic variability in their germ cells. This in turn should be translated into an increase of genetic variability inside the populations thus potentiating their evolutionary plasticity and evolvability.

## Introduction

Previous studies have shown in *Drosophila* and other organisms that the application of a heat shock stress may increase transcription levels of certain transposable elements also leading in some cases to their transposition “bursts”^1-4^. Most recently it has been also shown that the heat shock treatment to different *Drosophila* strains at pupal stage for several generations can induce mutations by transposon’s insertions^5^. Suggestions on the mechanism underlying such phenomenon come from the demonstration that functional alterations of the heat shock protein Hsp90, known as Hsp83 in *Drosophila*, causes the activation of various types of transposons in germ cells of *Drosophila*^6^, including a class of repetitive sequences called Stellate in male germ line, due to the involvement of the chaperone in the Piwi-interacting RNA (piRNA) biogenesis^7-9^. Such class of small interfering RNAs are located in the nuage, a specific electron-dense region around the nucleus of germ cells, where form ribonucleoprotein complexes called RNA-induced silencing complexes (RISCs). RISCs are in fact involved in a post-transcriptional mechanism that maintains transposable elements and repeated sequences in a repressed state^10-13^. Beside the piRNAs, two other classes of small RNAs, the small interfering RNAs (siRNAs) and microRNAs (miRNAs), are involved in post-transcriptional silencing in somatic cells^10,14,15^. All these classes of small RNAs perform their functions in post-transcriptional gene silence by RISCs. A fundamental component of RISCs is represented by proteins of the Argonaute (Ago) family: Ago1, Ago2 and Ago3 are respectively complexed with siRNAs, miRNAs and piRNAs. A suggestive point is the demonstration in cultured somatic cells of rat^16^, human^17,18^ and *Drosophila*^19^ that the ATP-dependent loading of both siRNAs and miRNAs duplexes to the Ago1 or Ago2 requires the aid of a Hsc70/Hsp90 chaperone machinery that seems to act with a coordinate mechanism similar to that involved in the activation of steroid hormone receptors upon the incorporation of the cognate ligand by the Hsp90 chaperone^20-22^. In *Drosophila*, it has been shown that for the loading of siRNAs and miRNAs to Ago1 and Ago2, the Hsc70/Hsp90 machinery also include other co-chaperones as Hop (Hsp70/Hsp90 organizing protein homolog)^23^, Hsc70-4 and Droj2 (DnaJ-like-2)^19^. This last factor belongs to the Hsp40 co-chaperones family which are essential in the Hsp70 cycle. These data together suggest that a similar chaperone machinery is probably required also for the loading of piRNAs to the Ago3 and seem to suggest a causal correlation between heat shock induced transposon activity and the functional destabilization of the Hsc70/Hsp90 chaperone machinery. We tested such hypothesis and here we show that heat shock activates transposons in germ cells by affecting the Hsc70/Hsp90 chaperone machinery with the heat-inducible Hsp70 chaperone as a major player in the piRNA pathway perturbation.

## Results and Discussion

First of all we wanted to analyze the impact of heat shock stress on transposons expression in our *Drosophila Ore-R* laboratory stock. Adult male and female flies were exposed to heavy heat shock (HHS: 37 °C for 1 h followed by 4 °C for 1 h, with the cycle repeated three times) and their germinal tissues were analyzed by qRT-PCR using oligonucleotides specific to different families of *Drosophila* retrotransposable elements. The results showed a significant increase of the transcripts of all transposons analyzed except copia in ovaries (Figure 1A).

**Figure 1.**
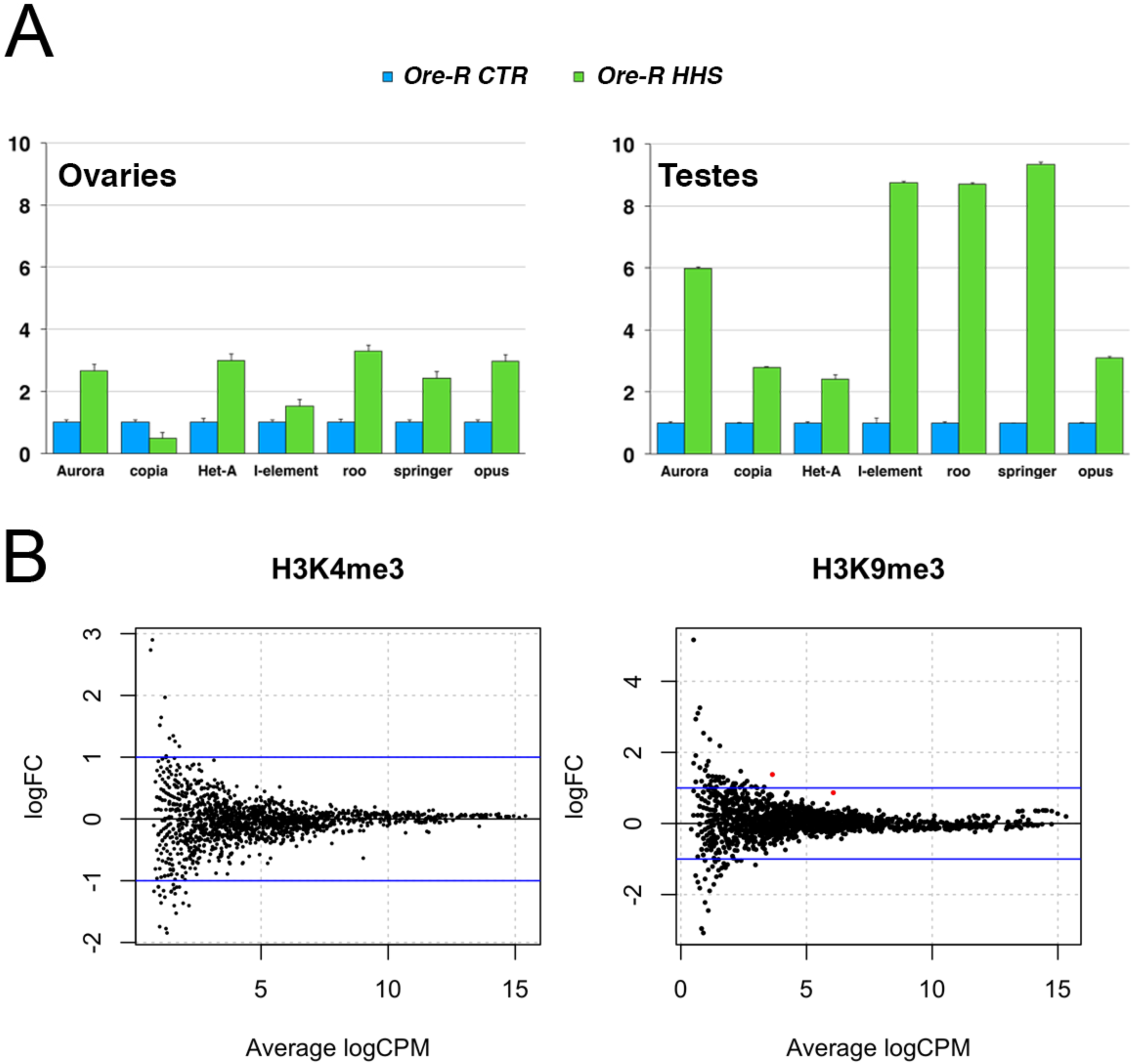
Heat shock treatment activates transposons by affecting their post-transcriptional silencing mechanism. (A) Transposable element expression profiles in ovaries and testes from control and heat-stressed flies analized two days after the heat treatment. Fold expression levels of transposon transcripts, determined by quantitative RT-PCR, are shown relative to Rp49 expression;(B) Plot showing the relationship between the log2-fold change between the two conditions (HHS and CTRL) versus the average log2-counts-per-million (CPM); the blue lines correspond to a fold change of two. Red color is used to highlight the statistically significant changes (with FDR< 0.05).

To test a possible effects of heat shock stress on transcriptional control of transposons we performed ChIP-seq experiments on ovaries extracts from heat treated and untreated females using antibodies against histone H3K9me3 and H3K4me3, two specific epigenetic markers for transcriptionally inactive and active chromatin respectively. We did not find significant differences of H3K4me3 marks on all the families of transposable elements before and after the treatment; in the H3K9me3 experiment, we found only 2 elements (FBti0060728 and FBti0063005) differentially enriched (FDR=0.009) with not great changes (log_2_FC=1.4 and 0.9 respectively) (Figure 1B). This result strongly suggests that the derepression of transposons after stress is mainly due to alterations of the post-transcriptional silencing mechanism. Since it has been shown that Hsp90 and Hop are involved in piRNA biogenesis^6,23^ we tested a possible involvement of Hsc70-4 and other co-chaperones such as Droj2 and dFKBP59 which are known key components of the molecular Hsc70-Hsp90 chaperone machinery^19,21^. To this end we firstly performed co-immunoprecipitation experiments on ovaries from non-stressed flies using a specific antibody against Ago3 and subsequent western blot analysis by specific antibodies. The results clearly demonstrated a binding interaction of Ago3 with Hsp90, Hop, Hsc70-4, and dFKBP59, a Hsp90-associated co-chaperone belonging to the class of immunophillins^24^ (Figure 2A). We performed also immunolocalization experiments and the results have shown that all above mentioned co-chaperones colocalize in ovaries to the nuage where are also located the other component of RISC such as Aubergine (Aub) and Ago3 (Figure 2B). We want to pointed out that we cannot check for the presence of Droj2 in both co-immunoprecipitation and immunofluorescence assays for the lack of specific antibody directed against this other key component of the Hsc70-Hsp90 complex. However, as reported below, we were able to analyze its functional involvement in piRNA biogenesis by RNAi silencing. To test whether the evidences for interactions and colocalizations in nuage of all such factors correspond to their functional requirement in transposon silencing, we analyzed the transposon expression profiles from ovaries in which we depleted Droj2, Hsc70-4 and dFKBP59 by *in vivo* RNAi using nanos-Gal4 driver (nosG4). We want to point out that both nanos-Gal4-mediated Droj2 and Hsc70-4 knockdown cause a complete ovarian and testes degeneration, as shown below, thus complicating the molecular analyses of TE transcripts; to bypass these developmental defects, we used the tub-Gal80^ts^ (tubG80^ts^) system to temporally control the expression of the dominant negative Hsc70-4^71S^ variant (DN-Hsc70-4) and Droj2 knockdown driven by nanos-Gal4. NosG4/ tubG80^ts^ >Droj2^RNAi^ and nosG tubG80^ts^ >DN-Hsc70-4 females were aged for 6 days at the permissive temperature (18°C) and then shifted to the restrictive temperature (29°C) for 5 days before dissecting the ovaries for RNA purification. As shown in Figure 3, we found, as already shown for Hsp90^6^ and Hop^23^, that the functional inhibition of Hsc70-4, dFKBP59 and Droj2 in gonads of non-stressed flies derepresses various classes of transposable elements in ovaries (Figure 3A) and activates the Stellate sequences (Ste) in testes as indicated by the presence of Ste crystalline aggregates (Figure 3B).

**Figure 2.**
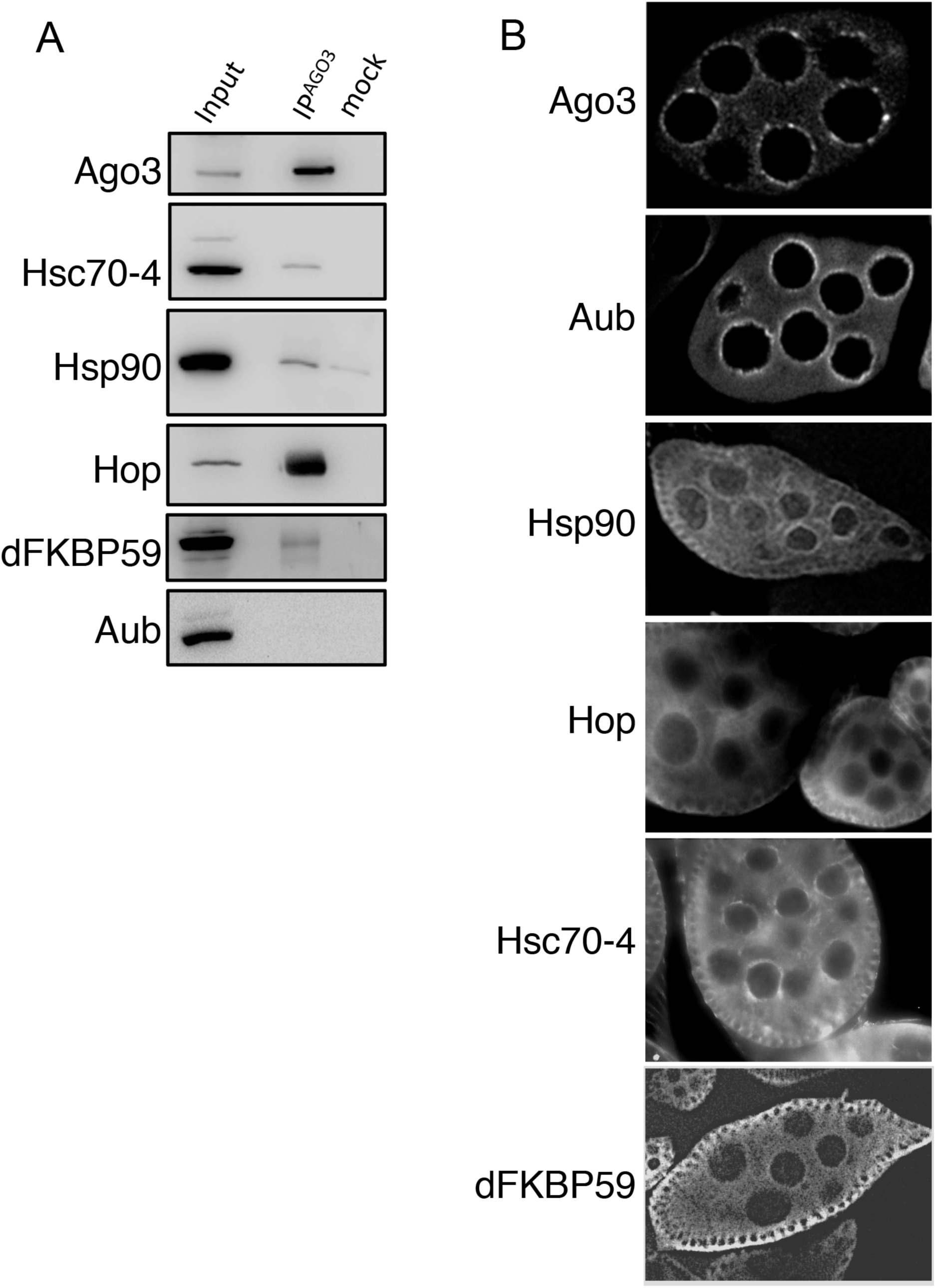
Interaction of Ago3 with components of the Hsc70-Hsp90 chaperone machinery. (A) Ago3 immunoprecipitates from wild-type ovaries subjected to western blotting analysis are probed with antibodies against the indicated co-chaperones. Co-immunoprecipitation experiments clearly show that all the protein tested with exception of Aubergine are direct interactors of Ago3; (B) immunofluorescence analysis confirms the co-localization of Ago3 with the same co-chaperones in the nuage. Note that, although not present in co-immunoprecipitation assay, Aubergine the other key player in piRNAs biogenesis, is also located in the nuage.

**Figure 3.**
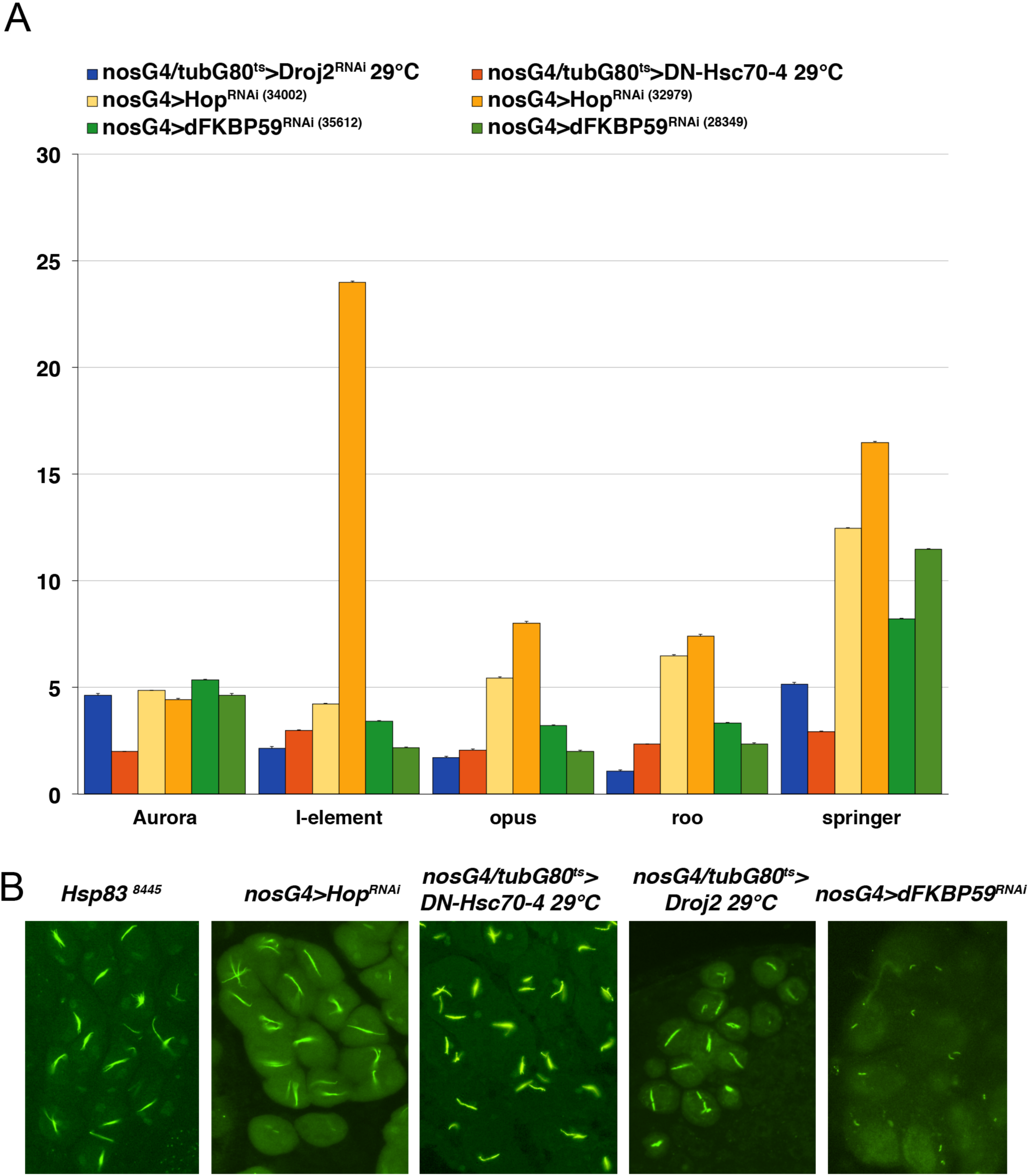
Components of the Hsp90-Hsp70 machinery are required for transposon silencing. (A) Diagram showing the activation of various transposons by silencing of co-chaperone genes Droj2, Hsc70-4, Hop and dFKBP59. Note that for both Hop and dFKBP59 two RNAi transgenic lines expressing by nos-G4 driver two different shRNAi constructs were tested while for Hsc70-4 the dominant negative variant DN-Hsc70-4 was tested. To temporally control the expression of DN-Hsc70-4 and Droj2 RNAi knockdown, the tub-Gal80^ts^ system was used; (B) the presence of crystals in co-chaperone depleted testes demonstrates that the same co-chaperones when silenced activate also the Stellate sequences as already know for the Hsp90 mutant allele *Hsp83*^*8445*^.

Together these data demonstrate that in normal conditions of temperature, transposons are largely repressed by the RNA interference mechanism in which the Hsc70/Hsp90 chaperone machinery plays an important role.

However, these data raise a critical point that needs to be addressed: although heat shock stress increases the protein levels of the above described chaperones, its effects on transposon expression are similar to those observed in the various chaperone mutants^6,23^. This apparent paradox could be resolved by assuming that heat shock stress partially displaces the chaperone complex, including also other factors such as Ago3, from its normal functions in piRNA biogenesis. This would establish an equivalence between a stress-induced functional shifting of the chaperone complex and the hypomorphic or RNAi mutations of its components. We supposed that a functional shifting of the chaperone machinery from the control of transposon activity after heat shock is probably caused by major heat shock protein Hsp70. In *Drosophila* Hsp70 is induced by heat shock and establishes an interaction with Hsp90 mediated by the co-chaperone Hop (Hsp70/Hsp90 organizing protein). So that, it appears reasonable to speculate that, after stress, the heat-induced Hsp70 establishes an interaction with the chaperone machinery, and perhaps other factors also involved in piRNA biogenesis, in forming new aggregates that should migrate outside the nuage for a probable degradation. The titration of the chaperone machinery and the other factors outside the nuage would affect piRNA biogenesis with the consequent activation of transposons.

To get insights about such possible interaction, we analyzed Hsp70 relationship with the Hsc70/Hsp90 machinery after heat shock in ovaries by immunostaining and immunoprecipitation experiments using specific antibodies. As shown in Figure 4A, after heat shock inducible Hsp70 localizes into nuage. Moreover, co-immunoprecipitation assay using a specific Hop antibody, shows that after 1h of recovery from heat shock, Hsp70 co-immunoprecipitates with Hop and Hsp90 in ovaries also together with another key player in the biogenesis of piRNA such as Ago3 (Figure 4B). Such results strongly suggest that all these factors physically interact in the nuage. To follow the fate of the interaction among these factors we firstly performed a localization analysis of Hsp90 and Hsp70 at different recovery times from heat stress on cytological preparations of ovaries. The results highlighted the delocalization of such proteins in cytoplasmic bodies formed outside of nuage after one and two days of recovery from heat shock (Figure 4C). This result acquires more significance considering that also the increase of transposon transcripts after heat shock occours after 2 days of recovery from treatment (see Figure 1A). At 3 days of recovery from stress, the effect begins to regress while at 7 days after stress the normal condition is restored. In addition, immunofluorescence experiments conducted on ovaries after 1 day from heat shock showed that also Ago3 and Hop localize in the same cytoplasmic bodies outside the nuage (Figure 4D). These results suggested that after 1 day of recovery from stress Hsp90, Hsp70, Hop and Ago3 localize in lysosomes probably for degradation. To test such suggestion, we analyzed ovaries preparations obtained after 1 day of recovery from heat shock by immunofluorescence experiments using the LysoTracker, a highly specific lysosomal marker and we observed the co-localization of all the factors with such marker (Figure 4D) thus confirming that such factors are in fact carried to lysosomes for degradation.

**Figure 4.**
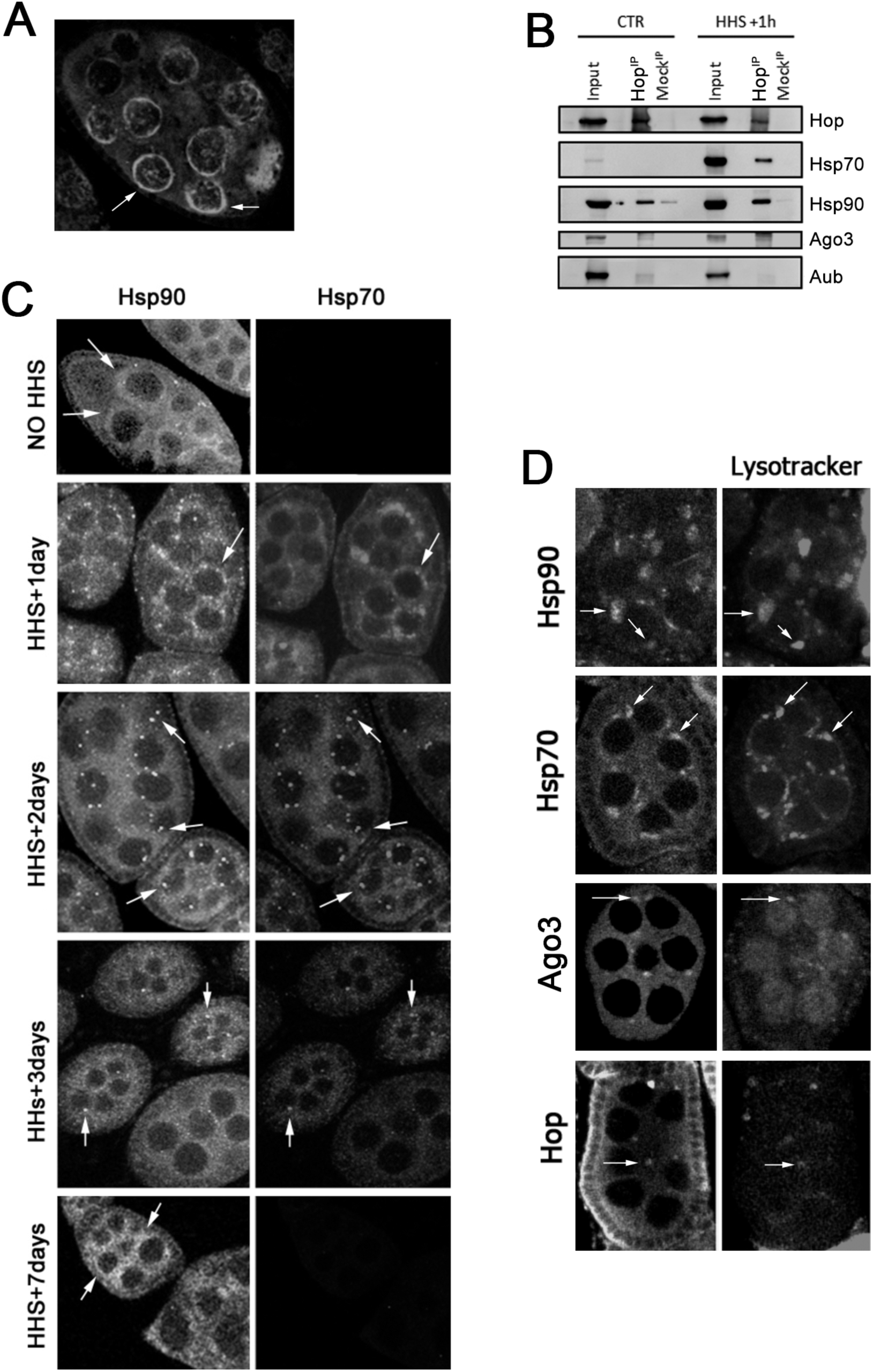
The heat shock induces an interaction of Hsp70 with Ago3 and the components of the chaperone machinery. (A) After 1hour from heat shock the induced Hsp70 localizes in the nuage and (B) establish an interaction with Ago3 and the Hsc70-Hsp90 machinery; (C) after 2 days from the heat shock Hsp70 interacts with Hsp90 in forming cytoplasmic bodies outside the nuage and after seven days Hsp90 appears again localized in the nuage while Hsp70 signals are not present;(D) importantly, Hsp90-Hsp70 cytoplasmic aggregates actually include also Ago3 and Hop that all together are finally delivered to lysosomes for degradation as shown by their co-localization with the specific lysosomal marker LysoTracker (see arrows for examples).

To understand whether cytoplasmic bodies observed in ovaries of stressed flies could be due to the dysfunction of Hsc70-Hsp90 machinery in piRNA pathway, we searched for the presence of these aggregates also in Hsp90 and Hop depleted ovaries in absence of stress (Figure 5A and B) and we found the presence of numerous and discrete cytoplasmic bodies where Ago3 localizes with Hop (Figure 5A) or Hsp90 (Figure 5B). Intriguingly, in both cases in which Hsp90 or Hop are silenced Hsp70 appears expressed and colocalized in cytoplasmic bodies. More strikingly we observed, in normal conditions of temperature, the expression of Hsp70 also in ovaries of flies in which other RISC components are silenced (Figure 5C). These data strongly suggest that the formation of cytoplasmic bodies observed after stress or in chaperone mutants could be related to the presence of Hsp70. We found a confirmation of such suggestion by using transgenic flies harboring an extra copy of *Hsp70Aa* gene under the control of UAS sequences^25^. When transgenic Hsp70Aa was ectopically expressed by nanos-Gal4, in normal conditions of temperatures, we found the formation of cytoplasmic bodies in ovaries (Figure 5D).

**Figure 5.**
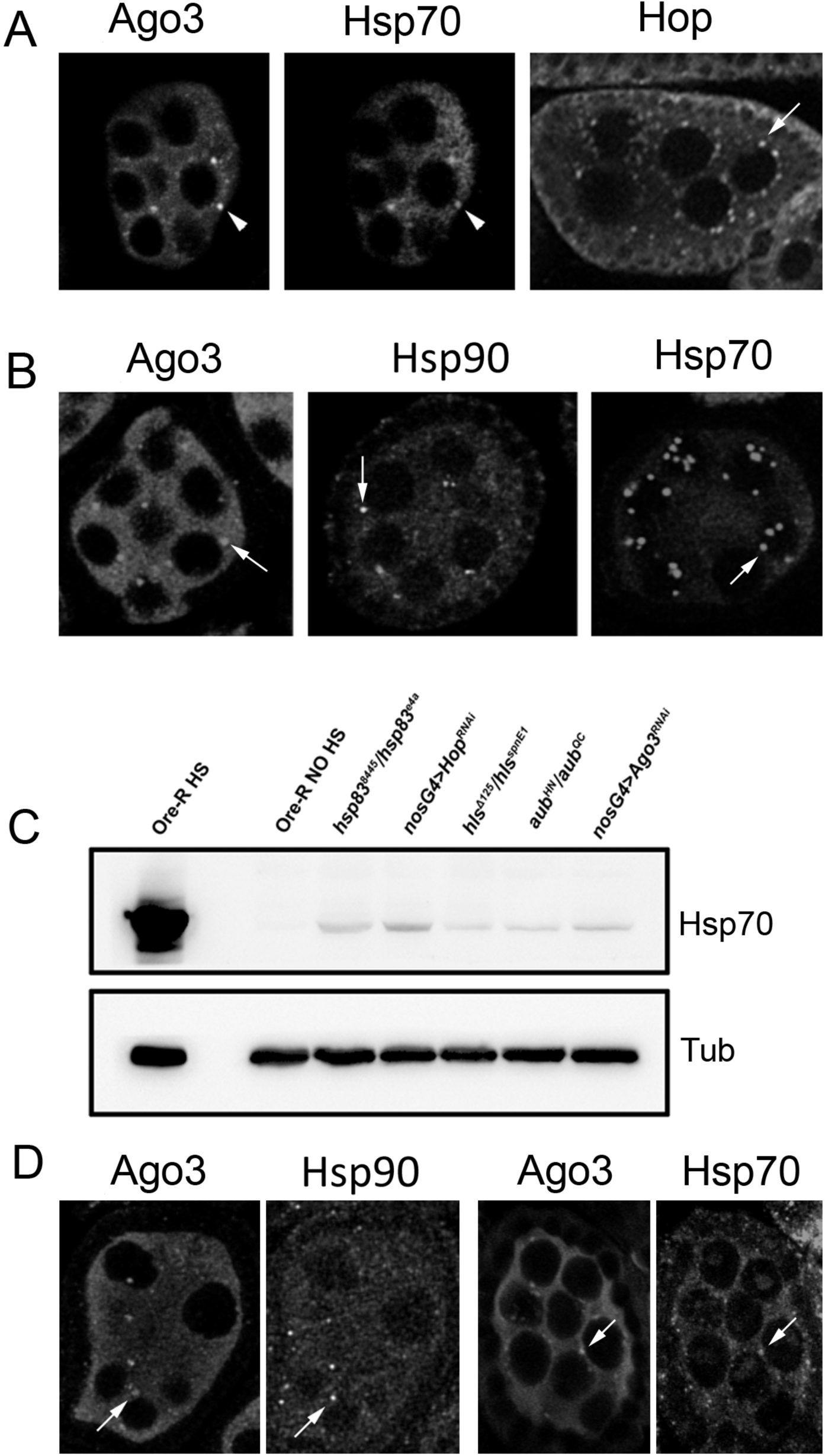
Functional inactivation of chaperones induces Hsp70 expression at normal temperatures along with its co-delocalization with Ago3 and the other chaperones in cytoplasmic bodies outside the nuage. (A) Ovaries from Hsp90 mutants showing cytoplasmic bodies outside the nuage (see arrows for examples) in which Ago3 co-localizes with Hop and Hsp70 (see arrowheads for an example); (B) in Hop depleted ovaries also Ago3 and Hsp90 accumulates in cytoplasmic bodies with Hsp70 (arrows for examples); (C) the quantitative reduction of chaperones as Hsp90 and Hop or other factors involved in piRNA biogenesis as Spindle-E, Ago3 and Aubergine induces the activation of Hsp70 chaperone at normal temperature conditions; (D) the expression of UAS-Hsp70 transgenic construct driven by nos-G4 at normal temperatures induces the formation of cytoplasmic bodies containing also Ago3 and Hsp90 (arrows for examples).

To assess the functional relevance of Hsp70 in transposons activation after stress, we wanted to investigate transposons activity in stressed flies carrying a complete deletion of Hsp70 genes cluster^26^. As shown in Figure 6, we found that in both testes and ovaries from heat shocked males and females lacking the Hsp70 cluster the increase of transposon transcripts is negligible compared to the observed increase of transposon transcripts observed in heat shocked wild type flies (see Figure 1A). This suggests a significant role of Hsp70 in stress induction of transposons expression. However, we want to point out that such conclusion seems to hold only for the results obtained in testes because we realized that, while the testes of heat shocked males appear normal (data not shown), the ovaries from heat shocked females lacking the Hsp70 cluster exhibit an early strong degeneration as confirmed by caspase-3 activation (Figure 7A and B). Moreover, we found such abnormal phenotype also in ovaries of heat shocked flies mutant for the heat shock factor (Hsf), a transcription factor necessary for Hsp70 activation after stress^27-30^ (Figure 7C-E). However, we found a significant increase of transposon transcripts after heat shock in ovaries of *hsf* mutant females expressing a *hsf*^*+t8*^ transgene that permits the Hsp70 induction^31^ (Figure 7F). This suggests that the expression of Hsp70 after stress is required not only for transposon expression but also to avoid ovary degeneration. A point of interest is that, differently from the other chaperones of the complex, the silencing of Hsc70-4 and Droj2 induce a strong ovarian (Figure 7G) and testes (Figure 7H) degeneration also in normal condition of temperature. These data show that Hsc70-4 plays a central role in ovaries and testes development at normal temperature while Hsp70 is necessary for the maintenance of ovaries after heat shock.

**Figure 6.**
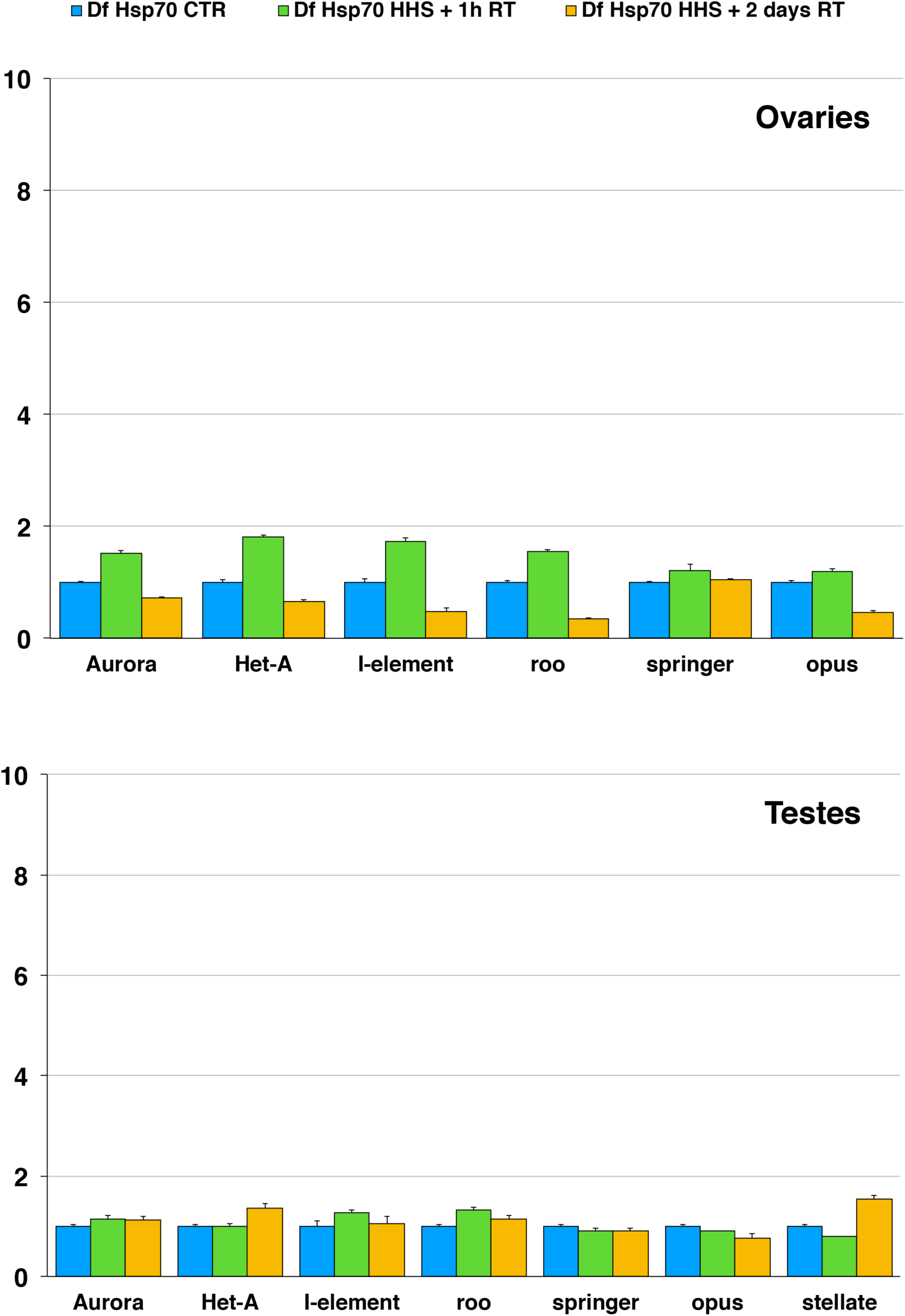
Heat shock has not significant effect on the amount of transposon transcripts in both ovaries and testes of flies carrying a complete deletion of Hsp70 genes cluster. In ovaries and testes the amount of transcripts of just some transposons is slightly increased after 1h of recovery from heat shock while for all transposons tested their transcripts results slightly decreased after 2 days of recovery. Also in testes the amount of transcripts is virtually not affected by heat shock at both 1h and 2 days from recovery.

**Figure 7.**
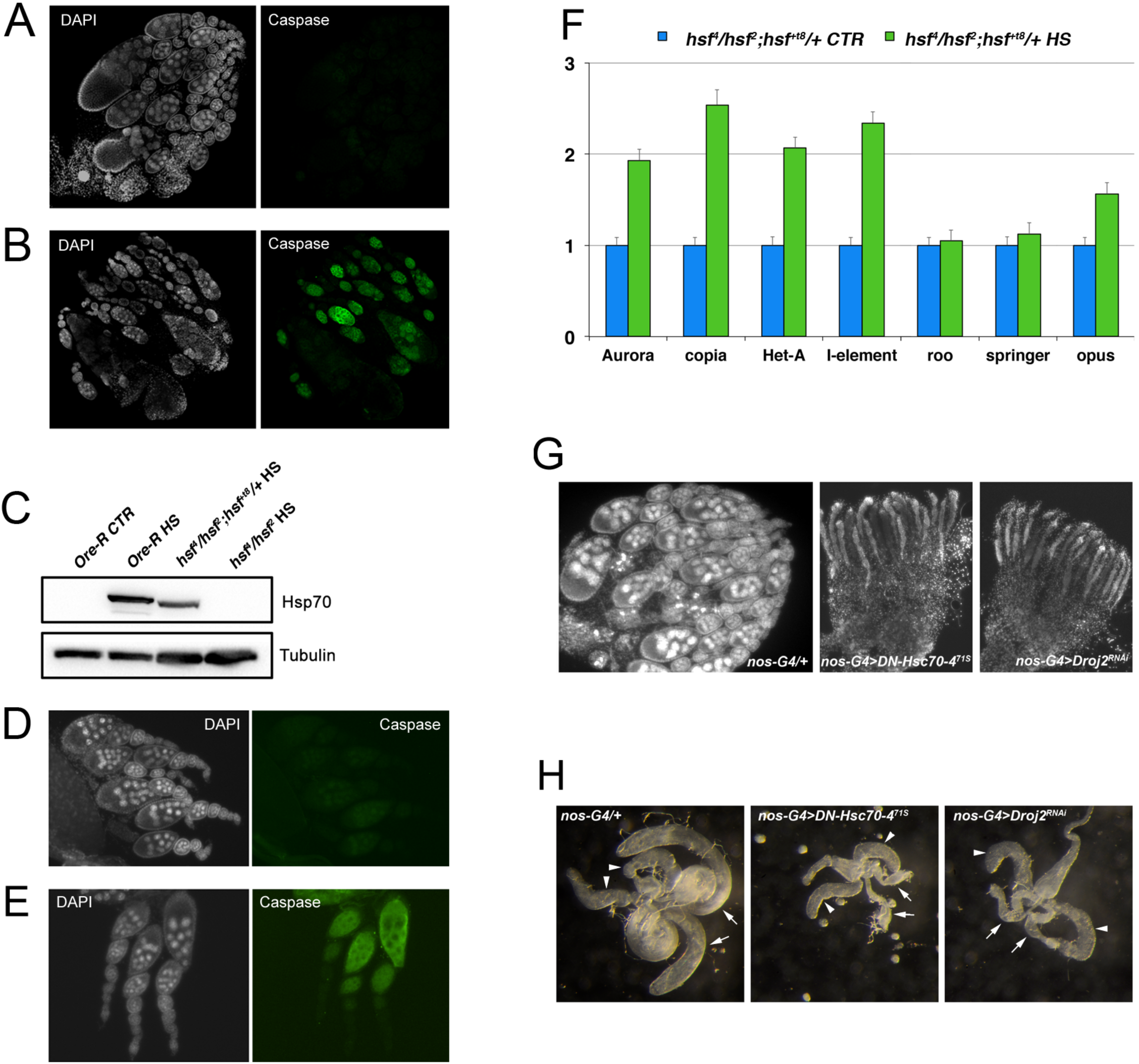
The heat shock stress produces a strong degeneration in ovaries from flies carrying a complete deficiency of the Hsp70 genes cluster. (A) The ovaries from Hsp70 deficient flies appear normal and without any signal of degeneration without heat shock; (B) after 24 hour from HHS treatment the ovaries contain *degenerating egg chambers* heavily stained by caspase-3, an enzyme involved in cell death; (C) western blot analysis shows that Hsp70 is not induced by heat shock in ovaries of females lacking the heat shock factor (hsf). Hsp70 can be instead induced by heat shock in *hsf* mutant females carrying a transgenic wild type copy (*hsf*^*+t8*^) of *hsf* gene; (D) also in ovaries from *hsf* trans-heterozygous mutants (*hsf*^*4*^ */hsf*^*2*^) no degeneration is observed without heat shock while (E) a strong degeneration after 24 hours from heat shock is clearly visible; (F) in ovaries from *hsf* mutant females carrying also a transgenic wild-type copy of *hsf*, an increased amount of transposons transcripts is detectable after heat shock; (G) Hsc70-4 and Droj2 mutant females show DAPI-stained ovaries with empty germaria arrested in stage 2; (H) Hsc70-4 and Droj2 mutant males show phase-contrast testes (arrows) that are abnormal comparing to those of wild type males (arrows) and lack spermatocytes (the arrowheads point the accessory glands).

As an alternative approach, we analyzed the amount of transposons transcripts in germ line of non-stressed transgenic flies expressing two different Hsp70 transgenes under UAS control and we found (Figure 8A) a significant activation of different transposons in ovaries while in testes we found a less pronounced increase of transcripts. However, in testes we have found numerous Ste crystalline aggregates in spermatocytes suggesting the activation of Stellate locus (Figure 8B-D) as confirmed by qRT-PCR on *Stellate* transcript (Figure 8A). These results demonstrate that ectopically expressed Hsp70 is capable to activate, also in normal condition, transposable elements and suggest that such chaperone is probably a main modulator of transposon activity after stress by interfering with the Hsc70/Hsp90 machinery.

**Figure 8.**
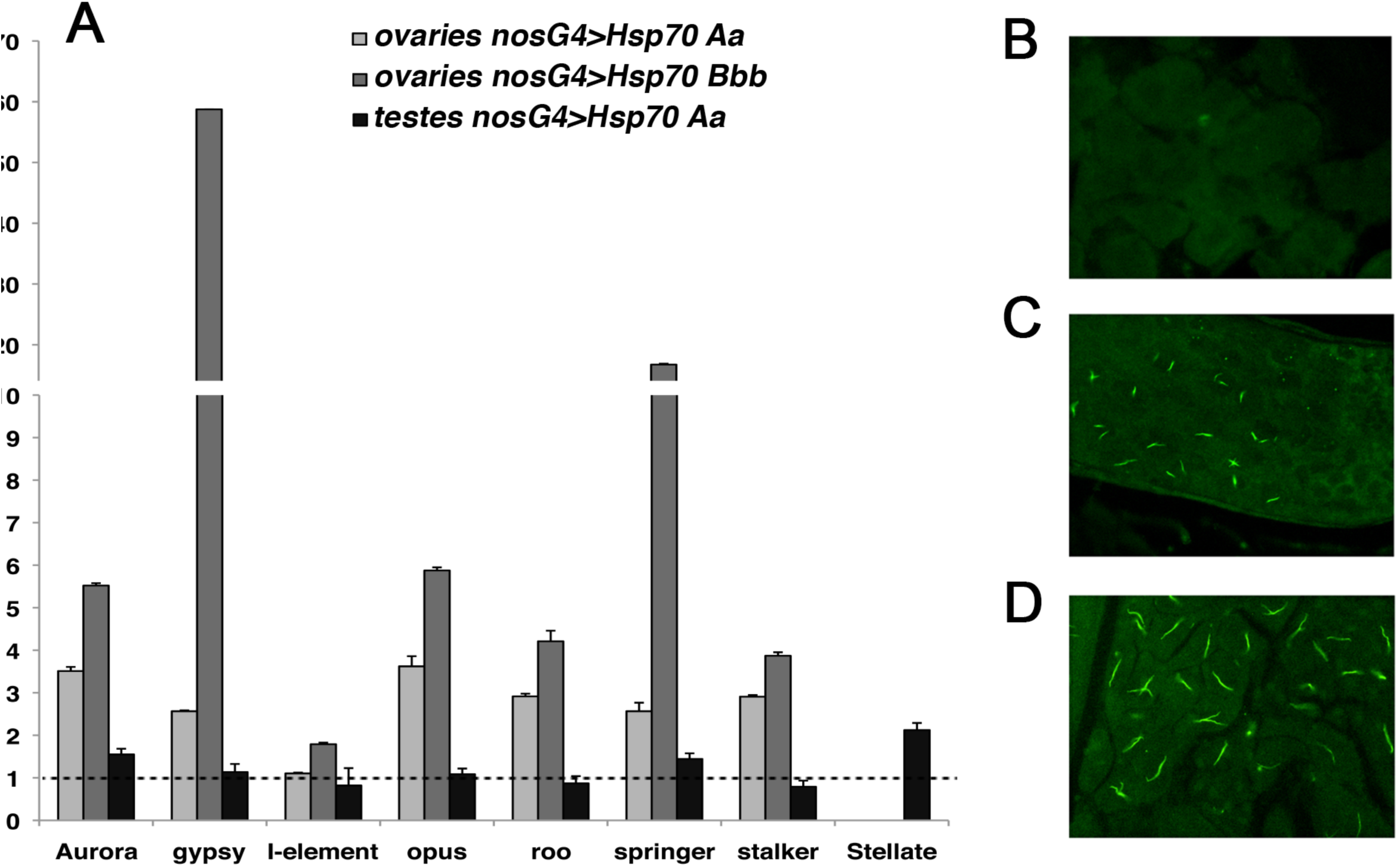
Hsp70 transgenes are able to induce both transposons and *Stellate (Ste)* activation in absence of stress; (A) qRT-PCR analysis shows a significant increase of transposon transcripts in ovaries from transgenic females expressing two genes of the Hsp70 genes cluster (Hsp70Aa and Hsp70Bbb). In testes the increase of transcripts is evident only for some of the transposons examined. The dashed line indicates the relative amount of TE transcripts in control ovaries; (B) testes from control males stained by a specific anti-Stellate antibody. No signals are evident; (C) testes from males carrying the *Hsp70Aa* and (D) *Hsp70Bbb* transgenes show the presence of crystalline aggregates after staining by anti-Ste antibody.

From all these results, we propose a model according to which the increase of transposons expression observed after heat shock is related to an active role of Hsp70 in delocalizing from the nuage to cytoplasmic bodies other stress affected factors that are involved in piRNA biogenesis. In conclusion, our results demonstrate that heat shock stress increases the expression of transposons mainly at post-transcriptional level by affecting piRNA biogenesis through the action of the inducible Hsp70 chaperone. Since several types of biotic and abiotic stress are able to trigger or reinforce the Hsp70 activity in both plants and animals^32^, we think that such mechanism contains relevant evolutionary implications. In presence of drastic environmental changes Hsp70 could play a key role not only in protecting the survival of individuals^26^ but also in increasing the frequency of mutations in their germ cells. This in turn should be translated into an increase of genetic variability inside the populations thus potentiating their environmental adaptability and evolvability^33^.

## Materials and Methods

### Drosophila strains

The *Drosophila* stocks used in this study were obtained from Bloomington Drosophila Stock Center (Indiana University, Bloomington, Indiana, USA) and are listed below.

Two hypomorfic allele of Hsp83 gene: *P{PZ}Hsp83*^*08445*^ *ry*^*506*^ (#11797) and *Hsp83*^*e4A*^ (Van Der Straeten et al., 1997); two indipendent Hop RNAi lines: *y*^*1*^ *sc* v*^*1*^; *P{TRiP.HMS00779}attP2* (#32979) and *y*^*1*^ *sc* v*^1^; *P{TRiP.HMS00965}attP2* (#34002); a line expressing a dominant negative allele of Hsc70-4, *w*^*126*^; *P{UAS-Hsc70-4.K*^*71S*^ *}G* (#5845); two indipendent dFKBP59 RNAi lines: *y*^*1*^ *sc* v*^*1*^; *P{TRiP.GL00453}attP2* (#35612) and *y*^*1*^ *v*^*1*^; *P{TRiP.JF02985}attP2*; a Droj2 RNAi line, *y*^*1*^ *sc* v*^*1*^; *P{TRiP.HMC04686}attP40* (#57382); a Ago3 RNAi line, *y*^*1*^ *sc* v*^*1*^; *P{TRiP.HMS00125}attP2*; a line expressing Hsp70Bbb under UAS control, *y*^*1*^ *w*^*67c23*^; *P{Mae-UAS.6.11}Hsp70Bbb*^*UY2168*^ (#7349); two hypomorphic alleles of Hsf gene: *cn*^*1*^ *bw*^*1*^ *Hsf*^*2*^ */CyO* (#5487) and *cn*^*1*^ *Hsf*^*4*^ *bw*^*1*^; *P{Hsf*^*+t8*^ *}1/TM3, Sb*^*1*^ *Ser*^*1*^; two mutant alleles of Aubergine gene: *aub*^*QC42*^ *cn*^*1*^ *bw*^*1*^ */CyO, l(2)DTS5131* (#4968) *and aub*^*HN2*^ *cn*^*1*^ *bw*^*1*^ */CyO* (#8517); two mutant alleles of Spindle E gene: *P{neoFRT}82B ry*^*506*^ *spn-E*^*hls-δ125*^ ^*e1*^ */TM3, ry* Sb*^*1*^ (#43638) and *ru*^*1*^ *st*^*1*^ *spn-E*^*1*^ *e*^*1*^ *ca*^*1*^ */TM3, Sb*^*1*^ *Ser*^*1*^ (#3327); a nanos-Gal4 driver, *P{UAS-Dcr-2.D}1, w*^*1118*^; *P{GAL4-nos.NGT}40* (#25751); a line encoding a temperature-sensitive form of the GAL80 repressor expressed under the control of the αTub84B promoter, *w*; P{tubP-GAL80*^*ts*^ *}2/TM2* (#7017). UAS-Hsp70Ab transgenic stock was provided by R. Meldrum Robertson (Department of Biology, Queen’s University, Kingston, Ontario K7L 3N6 Canada). Ore-R stock used here has been kept in our laboratory for many years. Flies were maintained and crossed at 24°C on standard cornmeal-sucrose-yeast-agar medium. To maximize the RNAi-mediated knockdown effect, flies were cultured at 29°C.

### Heavy HS (HHS) Treatment

5-7 days-old flies were treated at 37°C for 1 h followed by 4 °C for 1 h, with the cycle repeated three times.

### Co-Immunoprecipitation assays

Immunoprecipitation was performed according to Nishida et al., 2009^34^. Briefly, lysates from about 200 ovaries were prepared by homogenizing samples with a pestle grinder in 250 µl of Binding buffer (30 mM HEPES-KOH pH 7.3, 150 mM KOAc, 5 mM MgOAc, 5 mM DTT, 0.1% NP, 1×complete mini EDTA-free protease inhibitor cocktail). After centrifugation at 14 000 g for 1 min at 4°C, the supernatant was transferred to a new microcentrifuge tube and kept on ice. The pellet was then crushed again in 200 µl of Binding buffer. After centrifugation as above, the supernatant was combined with the first supernatant and kept on ice. The protein concentration of the lysates was adjusted each time to 4 mg/ml with Binding buffer. For immunoprecipitations rabbit anti-Ago3 (MAb Technologies) or mouse anti-Hop (a kind gift from M. Chinkers) antibodies were incubated with the lysates at 4°C for 4 hours in a rotating wheel. Dynabeads protein G (Invitrogen) were added and incubated overnight at 4°C. After 14 hours, the beads were washed two times in Binding buffer for 10 minutes and resuspended in an equal volume of SDS–PAGE sample buffer. After heating to 80°C for 10 min, protein samples were run on SDS–PAGE gels and processed for western blot analysis. 1% of the ovary lysate was run as an input sample.

### Western blot analyses

Protein extracts were fractionated by 10% SDS-PAGE and electroblotted onto Immobilion-P polyvinyl difluoride membranes (Bio-rad) in CAPS-based transfer buffer (10 mM CAPS pH 11, 10% methanol) in a semi-dry transfer apparatus (Amersham Biosciences). The membranes were blocked with 5% nonfat dry milk in Tris-buffered saline with Tween 20 (TBST) buffer (20 mM Tris pH 7.5, 150 mM NaCl, 0.1% Tween 20) and incubated with the following antibodies dissolved in TBST: mouse anti-Hsp90 antibody 1:2000 (a kind gift from R.Tanguay), rat anti-Hsp70 7FB antibody^35^ 1:3000 (provided by M. B. Evgen’ev), rabbit anti-Hop antibody 1:1000 and rabbit anti-dFKBP antibody 1:1000 (kindly provided by M.Chinkers), rabbit anti-Ago3 antibody 1:1000 (MAb Technologies), mouse anti-Aubergine antibody 1:2000 (a kind gift from M. Siomi), mouse anti-Hsc70 antibody 1:2000 (Sigma-Aldrich), mouse anti-αtubulin antibody 1:2000 (Sigma-Aldrich). The secondary antibodies were horseradish peroxidase-conjugated anti-mouse or anti-rabbit diluited 1:10000 in TBST. The Enhanced Chemiluminescence kit was purchased from GE Healthcare and the blot images were acquired with the ChemiDoc system (Bio-Rad Laboratories).

### ChIP-seq analysis

Approximately 300 dissected from 5-7 days-old females (heat-shocked and not-shocked) in 1X PBS (8.06 mM sodium phosphate, 1.94 mM potassium phosphate, 137 mM NaCl, 2.7mM KCl) were homogenization in A1 buffer (60 mM KCl, 15 mM NaCl, 15 mM Hepes pH 7.6, 4 mM MgCl_2_, 0.5% Triton X-100, 0.5 mM DTT, and 1× complete mini EDTA-free protease inhibitor cocktail) containing 1.8% formaldehyde and incubated for 15 min at room temperature (RT) in a rotating wheel. Cross-linking was stopped by adding glycine to 225 mM for 5 min at RT.

The homogenate was centrifuged for 5 min, 4000g at 4°C. The supernatant was discarded and the crude nuclei pellet was washed three times in A1 buffer and once in A2 buffer (140 mM NaCl, 15 mM Hepes pH 7.6, 1mM EDTA, 0.5 mM EGTA, 1% Triton X-100, 0.5 mM DTT, 0.1% sodium deoxycholate, and protease inhibitors) at 4°C. After the washes, nuclei were resuspended in A2 buffer in the presence of 0.1% SDS and 0.5% N-lauroylsarcosine, and incubated for 10 min in a rotating wheel at 4°C. After sonication (four pulses of 30 s with 1-min intervals, using a Hielscher Ultrasonic processor UP100H) and 10 min high-speed centrifugation, fragmented chromatin (DNA fragment size ranging from 200 bp to 500bp) was recovered in the supernatant. For each immunoprecipitation, 50 µg of chromatin was incubated in the presence of 10 µg of H3K4me3 (Active Motif) or H3K9me3 (Active Motif) antibodies (3 hours at 4°C in a rotating wheel). Then, 50 µl of dynabeads protein G (Invitrogen) was added and incubation was continued over night at 4°C. The supernatants were discarded and the beads were washed four times in A2 buffer supplemented with 0.05% SDS and twice in TE buffer (1 mM EDTA, 10 mM Tris pH 8)(each wash 5 min at 4°C). Chromatin was eluted from beads in two steps; first in 100 µl of 10 mM EDTA, 1% SDS, 50 mM Tris pH 8 at 65°C for 10 min, followed by centrifugation and recovery of the supernatant. Beads were re-extracted in 150 µl of TE, 0.67% SDS. The combined eluate (250 µl) was incubated 6 h at 65°C to reverse cross-links and treated with 50 µg /ml RNaseA for 15 min at 65°C and 500 µg /ml Proteinase K (Invitrogen) for 3 h at 65°C. Samples were phenol-chloroform extracted and ethanol precipitated. DNA was resuspended in 25 µl of water and sequenced by high-throughput Illumina technology.

### ChIP-seq analysis

Poor quality reads (more than 50% bases with quality scores < 28) have been filtered out. Any adaptor sequence or N-bases strings identified at read ends have been clipped by using Trimgalore software version 0.3.7 (https://github.com/FelixKrueger/TrimGalore). Trimmed reads shorter than 25 nucleotides have been discarded.

According to the strategy described in Rozhkov et al.^36^ the alignment of the reads has been made in two steps by using the Bowtie 2 software version 2.2.3^37^ Due to the repetitive nature of transposon sequences, reads were allowed to map to up to 10 different locations, with at maximum 2 mismatches on the D. melanogaster transposon sequences obtained from FlyBase version 6.09 (http://flybase.org/)^38^. Unaligned reads were then mapped onto genome. The dm6 release of the D. melanogaster genome was obtained from the University of California at Santa Cruz (UCSC) genome database (data not shown).

We have used 3 replicates for the H3K4me3 experiment and 2 replicates for the H3K9me3 one. For differential analysis referred to transposon sequences, the edgeR package^39^ has been used, keeping transposon locations with more than one count per million (CPM) in at least three samples for the H3K4me3 experiment and in at least two samples for the H3K9me3 experiment.

### Reverse transcription and Quantitative RT-PCR (qRT-PCR)

Total RNA purified according to the protocol supplied with Trizol Reagent (Invitrogen) was reverse transcribed using oligo dT and SuperScript reverse transcriptase III (Invitrogen). The qPCR reactions were carried out with QuantiFast SYBR Green PCR Kit (Qiagen) according to manufacturer’s protocol. For the quantification of transposon transcripts we used the 2^-δδCt^ method by comparing their amount to Rp49 transcript. qRT-PCR experiments were performed in three independent biological replicates each with three technical replicates. The primers used were:

Aurora F 5′-GAAGGAACTGAGCGTGTTCCA-3′

Aurora R 5′-CGTCTACCGCAACTAATGCAAA-3′

copia F 5′-TGGAGGTTGTGCCTCCACTT-3′

copia R 5′-CAATACCACGCTTAGTGGCATAAA-3′

Het-A F 5′-ACTGCTGAAGCTCGGATTCC-3′

Het-A R 5′-TGTAGCCGGATTCGTCATATTTC-3′

I element F 5′-CAATCACAACAACAAAATCC-3′

I element R 5′-GGTGTTGGTGTGGTTGGTTG-3′

roo F 5′-CGTCTGCAATGTACTGGCTCT-3′

roo R 5′-CGGCACTCCACTAACTTCTCC-3′

springer F 5′-CCATAACACCAGGGGCA-3′

springer R 5′-CGAGTGCTGGTCTGTCA-3′

opus F 5′-CGAGGAGTGGGGAGAGATTG-3′

opus R 5′-TGCGAAAATCTGCCTGAACC-3′

gypsy F 5′-CTTCACGTTCTGCGAGCGGTCT-3′

gypsy R 5′-CGCTCGAAGGTTACCAGGTAGGTTC-3′

stalker F 5′-TTATCAGGCTAGCCACATCTCTG-3′

stalker R 5′-TTGGCAGATATCACTTCTACCGATTC-3′

Stellate F 5′-GGCGATGAAAAGAAGTGG-3′

Stellate R 5′-CAGCGAGAAGAAGATGTC-3′

Rp49 F 5′-GCGCACCAAGCACTTCATC-3′

Rp49 R 5′-TTGGGCTTGCGCCATT-3′

#### Drosophila ovaries immunofluorescence

Ovaries were dissected in 1X PBS, fixed in 6% formaldehyde-methanol free in 1X PBS for 20 min, washed three times for 10 min in PBT (1X PBS/0.2% Triton X-100) and blocked in PBT-BSA (PBT plus 1% Bovin Serum Albumin) for 30 min. For LysoTracker staining, ovaries were dissected in PBS and the unfixed tissue was incubated in a 50 µM solution of LysoTracker (Invitrogen #L7528) in 1X PBS for 3 minutes. After rinsing with PBS, the tissues were fixed as described above. Ovaries were then incubated overnight at 4°C in PBT-BSA plus primary antibodies: rabbit anti-Hsp90 antibody 1:100 (Cell Signaling), rat anti-Hsp70 7FB antibody^37^ 1:300, mouse anti-Hsc70 antibody 1:200 (Sigma-Aldrich), rabbit anti-Hop antibody 1:400, rabbit anti-dFKBP59 antibody 1:400, mouse anti-Ago3 antibody 1:400 and mouse anti-Aubergine antibody 1:1000 (kindly provided by M. Siomi). Then the ovaries were washed three times for 10 min in PBT. Secondary antibodies were: FITC-conjugated goat anti-rat, Cy3-conjugated goat anti-rat, Alexa Fluor 555 goat anti-rabbit, Alexa Fluor 568 goat anti-mouse and Alexa Fluor 568 goat anti-rabbit (Life Technologies) (Life Technologies). The secondary antibodies were diluted in PBT/NGS (PBT plus 5% Normal Goat Serum NGS), and incubation was carried out 2h at room temperature. Ovaries were subsequently washed three times in PBT and incubated with TOTO-3 iodide (Invitrogen) for 30 minutes. After a wash in PBT the ovaries were separated into individual ovarioles and mounted in antifading medium (23,3 mg/ml of DABCO in 90% glycerol/10% 1X PBS) onto glass slides. Ovaries preparations were observed using a Zeiss Confocal Microscope.

#### Drosophilas testes immunofluorescence

Testes were dissected in TIT buffer (183 mM KCl, 47 mM NaCl, 10 mM Tris-HCl, 1 mM PMSF, 1 mM EDTA). After freezing in liquid nitrogen the tissues were fixed in cold methanol for 5 minutes and incubated in acetone for 1 minute. Then the testes were washed in 1X PBS for 10 minutes and incubated overnight with rabbit anti-Stellate antibody 1:50 (kindly provided by M. P. Bozzetti). After three wash in 1X PBS, testes were incubated for 1 h at room temperature with secondary antibody: Alexa Fluor 488 goat anti-rabbit (Life Technologies). The preparation was washed three times in 1X PBS, incubated with DAPI for 4 minutes and mounted in antifading medium onto glass slides.

## Acknowledgments

Financial support for this research was provided by the Epigenomics Flagship Project EpiGen, the Italian Ministry of Education and Research, National Research Council. We thank Bloomington *Drosophila* Stock Center for flies′ stocks and M. B. Evgen’ev, M. Siomi, M. Chinkers, R. Tanguay,

M. P. Bozzetti for antibodies.

## References

1. Strand DJ, McDonald JF. (1985). Copia is transcriptionally responsive to environmental stress. Nucleic Acids Res. 13(12):4401–10

2. Junakovic N, Di Franco C, Barsanti P, Palumbo G. (1986). Transposition of copia-like nomadic elements can be induced by heat shock. J Mol Evol. 24(1-2):89–93

3. Ratner VA, Zabanov SA, Kolesnikova OV, Vasilyeva LA. (1992). Induction of the mobile genetic element Dm-412 transpositions in the Drosophila genome by heat shock treatment. Proc Natl Acad Sci U S A. 89(12):5650–4

4. Arnault C, Dufournel I. (1994). Genome and stresses: reactions against aggressions, behavior of transposable elements. Genetica. 93(1-3):149–60

5. Fanti L, Piacentini L, Cappucci U, Casale AM, Pimpinelli S. (2017). Canalization by Selection of de Novo Induced Mutations. Genetics. 206(4):1995–2006

6. Specchia V, Piacentini L, Tritto P, Fanti L, D′Alessandro R, Palumbo G, Pimpinelli S, Bozzetti MP. (2010). Hsp90 prevents phenotypic variation by suppressing the mutagenic activity of transposons. Nature. 463(7281):662–5

7. Miyoshi T, Takeuchi A, Siomi H, Siomi M.C. (2010). A direct role for Hsp90 in pre-RISC formation in Drosophila. Nat. Struct. Mol. Biol. 17(8): 1024–1026

8. Luteijn MJ, Ketting RF. (2013). PIWI-interacting RNAs: from generation to transgenerational epigenetics. Nat Rev Genet 14(8): 523-534

9. Iwasaki YW, Siomi MC, Siomi H. (2015). PIWI-Interacting RNA: Its Biogenesis and Functions. Annu Rev Biochem. 84:405–33

10. Ghildiyal M, Zamore PD. (2009). Small silencing RNAs: an expanding universe. Nat Rev Genet 10(2): 94–108

11. Johnston M, Geoffroy M, Sobala A, Hay R, Hutvagner G. (2010). HSP90 protein stabilizes unloaded Argonaute complexes and microscopic P-bodies in human cells. Mol Biol Cell 21(9): 1462–1469

12. Iki T, Yoshikawa M, Nishikiori M, Jaudal M, Matsumoto-Yokoyama E, Mitsuhara I, Meshi T, Ishikawa M. (2010). In vitro assembly of plant RNA-induced silencing complexes facilitated by molecular chaperone HSP90. Mol Cell 39(2): 282–291

13. Gangaraju VK, Yin H, Weiner MM, Wang J, Huang XA, Lin H. (2011). Drosophila Piwi functions in Hsp90-mediated suppression of phenotypic variation. Nat Genet. 43(2):153–8

14. Carthew RW, Sontheimer EJ. (2009). Origins and Mechanisms of miRNAs and siRNAs. Cell. 136(4):642–55

15. Kim VN, Han J, Siomi MC. (2009). Biogenesis of small RNAs in animals. Nat Rev Mol Cell Biol. 10(2):126–39

16. Tahbaz N, Carmichael JB, and Hobman TC. (2001). GERp95 belongs to a family of signal-transducing proteins and requires Hsp90 activity for stability and Golgi localization. J. Biol. Chem. 276:43294–43299

17. Höck J, Weinmann L, Ender C, Rüdel S, Kremmer E, Raabe M, Urlaub H, Meister G. (2007). Proteomic and functional analysis of Argonaute-containing mRNA-protein complexes in human cells. EMBO Rep. 8(11):1052–60

18. Landthaler M, Gaidatzis D, Rothballer A, Chen PY, Soll SJ, Dinic L, Ojo T, Hafner M, Zavolan M, Tuschl T. (2008). Molecular characterization of human Argonaute-containing ribonucleoprotein complexes and their bound target mRNAs. RNA. 14(12):2580–96

19. Iwasaki S, Kobayashi M, Yoda M, Sakaguchi Y, Katsuma S, Suzuki T, Tomari Y. (2010). Hsc70/Hsp90 chaperone machinery mediates ATP-dependent RISC loading of small RNA duplexes. Mol. Cell 39(2): 292–299

20. Kirschke E, Goswami D, Southworth D, Griffin PR, Agard DA. (2014). Glucocorticoid receptor function regulated by coordinated action of the Hsp90 and Hsp70 chaperone cycles. Cell. 157(7):1685–97

21. Li J, Soroka J, Buchner J. (2012) The Hsp90 chaperone machinery: conformational dynamics and regulation by co-chaperones. Biochim Biophys Acta. 1823(3):624–35

22. Tsuboyama K, Tadakuma H, Tomari Y. (2018). Conformational Activation of Argonaute by Distinct yet Coordinated Actions of the Hsp70 and Hsp90 Chaperone Systems. Mol Cell. 70(4):722–729

23. Karam JA, Parikh RY, Nayak D, Rosenkranz D, Gangaraju VK. (2017). Co-chaperone Hsp70/Hsp90-organizing protein (Hop) is required for transposon silencing and Piwi-interacting RNA (piRNA) biogenesis. J Biol Chem. 292(15):6039–6046

24. Zaffran S. (2000). Molecular cloning and embryonic expression of dFKBP59, a novel Drosophila FK506-binding protein. Gene. 246(1-2):103–9

25. Xiao C, Mileva-Seitz V, Seroude L, Robertson RM. (2007). Targeting HSP70 to motoneurons protects locomotor activity from hyperthermia in Drosophila. Dev Neurobiol. 67(4):438–55

26. Gong WJ, Golic KG. (2004). Genomic deletions of the Drosophila melanogaster Hsp70 genes. Genetics. 168(3):1467–76

27. Lis J, Wu C. (1993) Protein traffic on the heat shock promoter: parking, stalling, and trucking along. Cell. 74(1):1–4

28. Morimoto RI. (1993). Cells in stress: transcriptional activation of heat shock genes. Science. 259(5100):1409–10

29. Voellmy R. (1994). Transduction of the stress signal and mechanisms of transcriptional regulation of heat shock/stress protein gene expression in higher eukaryotes. Crit Rev Eukaryot Gene Expr. 4(4):357–401

30. Wu C. (1995). Heat shock transcription factors: structure and regulation. Annu Rev Cell Dev Biol. 11:441–69

31. Jedlicka P, Mortin MA, Wu C. (1997). Multiple functions of Drosophila heat shock transcription factor in vivo. EMBO J. 16(9):2452–62.

32. Yu A, Li P, Tang T, Wang J, Chen Y3, Liu L. (2015). Roles of Hsp70s in Stress Responses of Microorganisms, Plants, and Animals. Biomed Res Int. 2015: 510319

33. Piacentini L, Fanti L, Specchia V, Bozzetti MP, Berloco M, Palumbo G, Pimpinelli S. (2014). Transposons, environmental changes, and heritable induced phenotypic variability. Chromosoma. 123(4):345–54

34. Nishida KM, Okada TN, Kawamura T, Mituyama T, Kawamura Y, Inagaki S, Huang H, Chen D, Kodama T, Siomi H, Siomi MC. (2009). Functional involvement of Tudor and dPRMT5 in the piRNA processing pathway in Drosophila germlines. EMBO J. 28(24):3820–31

35. Velazquez JM, Lindquist S. (1984). Hsp70: nuclear concentration during environmental stress and cytoplasmic storage during recovery. Cell. 36:655–662

36. Rozhkov NV, Hammell M, Hannon GJ. (2013). Multiple roles for Piwi in silencing Drosophila transposons. Genes Dev 27(4):400–12

37. Langmead B, Salzberg SL. (2012). Fast gapped-read alignment with Bowtie 2. Nat Methods. 9(4):357–9

38. Attrill H, Falls K, Goodman JL, Millburn GH, Antonazzo G, Rey AJ, Marygold SJ; FlyBase Consortium. (2016). FlyBase: establishing a Gene Group resource for Drosophila melanogaster. Nucleic Acids Res. 44(D1):D786–92

39. Robinson MD, McCarthy DJ, Smyth GK. (2010). edgeR: a Bioconductor package for differential expression analysis of digital gene expression data. Bioinformatics. 26(1):139–4

